# Rapid, dose-dependent and efficient inactivation of surface dried SARS-CoV-2 by 254 nm UV-C irradiation

**DOI:** 10.1101/2020.09.22.308098

**Authors:** Natalia Ruetalo, Ramona Businger, Michael Schindler

## Abstract

**Background:** The SARS-CoV-2 pandemic urges for cheap, reliable, and rapid technologies for disinfection and decontamination. One frequently proposed method is UV-C irradiation. However, UV-C doses necessary to achieve inactivation of high-titer SARS-CoV-2 are poorly defined.

**Methods:** Using a box and two handheld systems designed to decontaminate objects and surfaces we evaluated the efficacy of 254 nm UV-C treatment to inactivate surface dried SARS-CoV-2.

**Results:** Drying for two hours did not have a major impact on the infectivity of SARS-CoV-2, indicating that exhaled virus in droplets or aerosols stays infectious on surfaces at least for a certain amount of time. Short exposure of high titer surface dried virus (3-5*10^6 IU/ml) with UV-C light (16 mJ/cm^2^) resulted in a total inactivation of SARS-CoV-2. Dose-dependency experiments revealed that 3.5 mJ/cm^2^ were still effective to achieve a > 6-log reduction in viral titers whereas 1.75 mJ/cm^2^ lowered infectivity only by one order of magnitude.

**Conclusions:** Our results demonstrate that SARS-CoV-2 is rapidly inactivated by relatively low doses of UV-C irradiation. Furthermore, the data reveal that the relationship between UV-C dose and log-viral titer reduction of surface residing SARS-CoV-2 is non-linear. In the context of UV-C-based technologies used to disinfect surfaces, our findings emphasize the necessity to assure sufficient and complete exposure of all relevant areas by integrated UV-C doses of at least 3.5 mJ/cm^2^ at 254 nm. Altogether, UV-C treatment is an effective non-chemical possibility to decontaminate surfaces from high-titer infectious SARS-CoV-2.

## Introduction

SARS-CoV-2 has spread globally and there is an urgent need for rapid, highly efficient, environmentally friendly, and non-chemical disinfection procedures. Application of UV-C light is an established technology for decontamination of surfaces and aerosols (1–3). This procedure has proven effective to inactivate SARS-CoV-1 (4–6), several other enveloped and non-enveloped viruses as well as bacteria (7). UV-C-based disinfection could be applied in operating rooms and healthcare facilities but also prove useful in the business sector, where there is the necessity to sterilize surfaces being frequently encountered by multiple different individuals. Some examples also discussed in the context of public health are escalators, public transportation, rental cars, door handles and waiting rooms. Recently, it has also been shown that SARS-CoV-2 is sensitive to inactivation by UV-C irradiation (8–12). However, the aforementioned studies used high UV-C doses from 108 mJ/cm^2^ to more than 1 J/cm^2^ at exposure times from 50 seconds to several minutes necessary for total inactivation of SARS-CoV-2 (10–12). These parameters are in a range complicating efficient application of UV-based methods to be employed for large-scale decontamination of surfaces and aerosols. Others used innovative 222 nm or 280 nm UV-C LED technologies (8, 9) which are not yet implemented in most established 254 nm UV-C-based decontamination devices and needed relatively high doses of UV-C irradiation for inactivation, too. Another recent study by the Boston University established 254 nm UV-C dose-dependency inactivation kinetics of SARS-CoV-2 and reported doses necessary for complete sterilization of dry and wet virus preparations between 4 s and 9 s at 0.85 mW/cm^2^ in a test box (13). While this data is promising, a limitation was the study design in a test box and relatively low viral titers used, just allowing to conclude 2- to 3-log titer reductions by the treatment. Overall, the exact knowledge about dose-dependent inactivation kinetics is essential to design UV-C-based decontamination procedures that allow firm disinfection of SARS-CoV-2.

We hence conducted an approach simulating the inactivation of dried surface residing high-titer infectious SARS-CoV-2 by two mobile handheld UV-C emitting devices and an UV-C box designed to decontaminate medium-size objects. We asked the question of whether short exposure of SARS-CoV-2 to UV-C irradiation is sufficient to reduce viral infectivity and which UV-C doses are necessary to achieve an at least 6-log reduction in viral titers.

## Material and Methods

### Cell culture

Caco-2 (Human Colorectal adenocarcinoma) cells were cultured at 37 °C with 5% CO_2_ in DMEM (Dulbecco’s Modified Eagle Medium) containing 10% FCS, with 2 mM l-glutamine, 100 μg/ml penicillin-streptomycin and 1% NEAA (Non-Essential Amino Acid).

### Viruses

The recombinant SARS-CoV-2 expressing mNeonGreen (icSARS-CoV-2-mNG) (14) was obtained from the World Reference Center for Emerging Viruses and Arboviruses (WRCEVA) at the UTMB (University of Texas Medical Branch). To generate icSARS-CoV-2-mNG stocks, 200,000 Caco-2 cells were infected with 50 μl of virus stock in a 6-well plate, the supernatant was harvested 48 hpi, centrifuged, and stored at −80°C. For MOI (Multiplicity of infection) determination, a titration using serial dilutions of the virus stock was conducted. The number of infectious virus particles per ml was calculated as the (MOI × cell number)/(infection volume), where MOI = −ln(1 – infection rate).

### UV-C light inactivation treatment

35 μl of virus stock, corresponding to ~4-6*10^6^ infectious units (IU) of icSARS-CoV-2-mNG were spotted (in triplicates) in 6-well plates and dried for two hours at RT. This setup was chosen to mimic the situation in which an infected person exhales droplets that dry on surfaces and potentially stay infectious and hazardous over a prolonged period of time. 6-well plates spotted with dried virus were treated with UV-C-light (254 nm) using the Soluva^®^ pro UV Disinfection Chamber (Heraeus) for 60 seconds or the Soluva^®^ Zone HP Disinfection Handheld (Heraeus) for 2 seconds in a fix regime at 5 and 20 cm plate distance. In addition, a moving regime using a slow (3.75 cm/s) and fast (12 cm/s) speed at 20 cm distance was tested. Additionally, we employed a 2^nd^ generation Disinfection Handheld Soluva^®^ Zone H (Heraeus) which is less powerful than the Soluva^®^ pro UV but works autonomously with a rechargeable battery. See the spectrum of UV-C lamps employed in these devices in Supplemental Image 1. The lower UV-C intensity emitted by this device allowed us to perform a dose-dependency experiment exposing dried virus with different UV-C intensities.

The time dependent UV-C intensity emitted by the Soluva^®^ Zone H at various distances is detailed and depicted in Supplemental Image 2. UV exposure was carried out after 10 minutes of pre-heating the device at a distance of 50 cm for 20 s, 10 s, 5 s, 2.5 s, 20 s + 97 % UV-filter, 10 s + 97 % UV-filter corresponding to 14 mJ/cm^2^, 7 mJ/cm^2^, 3.5 mJ/cm^2^, 1.75 mJ/cm^2^, 0.42 mJ/cm^2^ and 0.21 mJ/cm^2^. These values are based on an on-site and parallel measurement of UV-C intensity emitted by the device via an UV-C dosimeter (Dr. Gröbel UV electronic GmbH), which corresponds to 0.7 mJ/cm^2^ when the UV-C light is applied at 50 cm distance, which fits quite well to the previously company measured value of 0.84 mJ/cm^2^ (Supplemental Image 2). As control, 6-well plates were spotted with the virus and dried, but not UV-treated. After UV-treatment, the spotted virus was reconstituted using 1 ml of infection media (culture media with 5% FCS) and viral titers determined as explained below. As additional control, 35 μl of the original virus stock were diluted to 1 ml with infection media and used as virus stock infection control. All UV-treatments were done at RT.

### Evaluation of UV-treatment

For infection experiments and titer determination, 1 ×10^4^ Caco-2 cells/well were seeded in 96-well plates the day before infection. Cells were incubated with the SARS-CoV-2 strain icSARS-CoV-2-mNG at a MOI=1.1 (stock) or the UV-treated and reconstituted virus in serial two-fold dilutions from 1:200 up to 1:51200 and in one experiment up to 1:102400. 48 hpi cells were fixed with 2% PFA (Paraformaldehyde) and stained with Hoechst33342 (1 μg/ml final concentration) for 10 minutes at 37°C. The staining solution was removed and exchanged for PBS (Phosphate-buffered saline). For quantification of infection rates, images were taken with the Cytation3 (Biotek) and Hoechst+ and mNG+ cells were automatically counted by the Gen5 Software (Biotek). Viral titers (number of infectious virus particles per ml) were calculated as the (MOI × cell number)/(infection volume), where MOI = −ln(1 – infection rate). Infection rates lower than 0.01 were used as a cutoff and set to 0 in order to avoid false positive calculations.

### Software and statistical analysis

Experiments were repeated two to four times each using duplicate or triplicate infections. GraphPad Prism 8.0 was used for statistical analyses and to generate graphs, as well as CorelDrawX7. Other software used included Gen5 v.3.10.

## Results

### Inactivation of high-titer SARS-CoV-2 by UV-C treatment

We set up an experimental approach to evaluate the effect of UV-C treatment on the infectivity of SARS-CoV-2. Simulating the situation that exhaled droplets or aerosols from infected individuals contaminate surfaces, we produced a high-titer SARS-CoV-2 infectious stock and dried 35μl of this stock corresponding to ~4-6*10^6 IU/ml in each well of a 6-well plate. The plates were then either non-treated or exposed to five UV-C regimens at 254 nm (Fig. 1a). These include inactivation for 60 s in a box designed to disinfect medium-size objects, 2 s exposure at 5 cm or 20 cm distance with a handheld UV-C disinfection device and an approach simulating decontamination of surfaces via the handheld UV-C device (Zone HP). For this, we performed slow and fast-moving at a distance of ~20 cm, with “slow” corresponding to a speed of ~3.75 cm/s (supplemental movie 1) and “fast” at ~12 cm/s (supplemental movie 2). UV-C irradiance (254 nm) in the box with an exposure time of 60 seconds corresponds to an irradiation dose of 600 mJ/cm^2^; for the handheld (HH) at 5 cm the UV-C dose at two second irradiation time is 80 mJ/cm^2^ and at 20 cm is 16 mJ/cm^2^. From the speed of the “slow” and “fast” moving regimens we calculate a UV-C dose of 2.13 mJ/cm^2^ (slow) and 0.66 mJ/cm^2^ (fast), assuming a focused intensity beam. However, taking into consideration the UV-C light distribution underneath the handheld device the integrated UV-C dose accumulates to 20 mJ/cm^2^ for the fast regimen.

**Figure 1.**
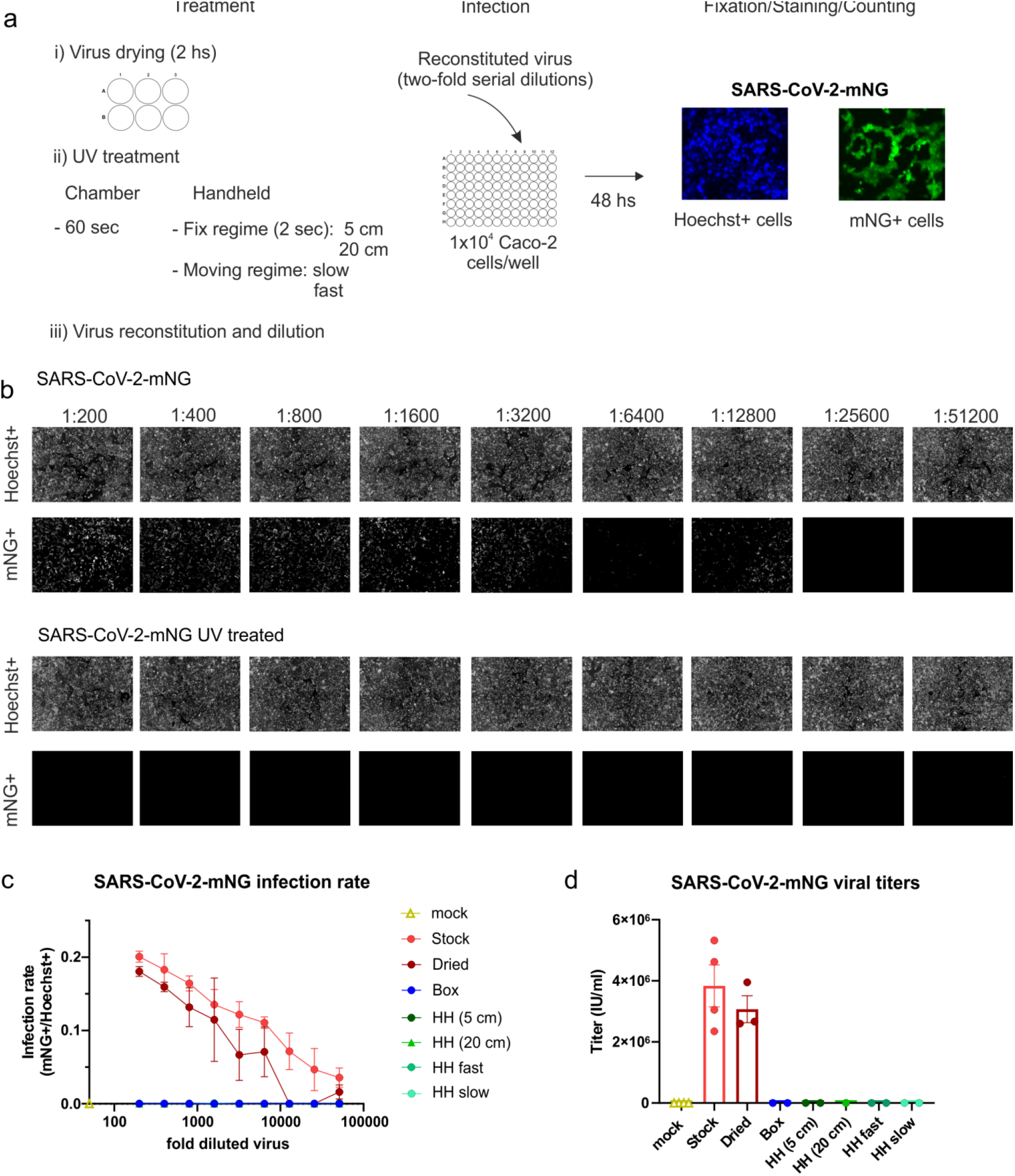
Inactivation of SARS-CoV-2 by UV-C light treatment. (a) Experimental layout of the different UV-treatments and the infection assay employed using the green-fluorescent virus SARS-CoV-2.mNG. (b) Primary data showing the results of the infection assay using the non-treated stock virus as a positive control and the UV-treated virus (HH, fast-moving regime). In the upper row, the total amount of cells for each well of the two-fold serial dilution of virus is shown as Hoechst+. In the lower, infected cells are visualized indicated as mNG+ cells. (c) Infection rate curves for UV-irradiated SARS-CoV-2-mNG using different UV-treatments. The graph shows the infection rate at each two-fold serial dilution, calculated as the number of infected cells (mNG+) over the total number of cells (Hoechst+) for the non-treated viral stock (n=4), dried viral stock (n=3), and dried and UV-irradiated virus using five different UV-treatments (n=2). Data are presented as mean +/- SEM of the number of biological replicates indicated above. (d) SARS-CoV-2-mNG viral titers after UV-treatment. The graph shows the viral titers calculated in IU/mL for the mock-infected, non-treated, and dried stock as well as the dried and UV-irradiated virus under the different treatments. The number of biological replicates (n=2-4) is directly plotted and indicated in 1c. Data are presented as mean +/- SEM.

Subsequently, dried virus was reconstituted with 1 mL infection media and used to inoculate naïve Caco-2 cells at serial dilutions to calculate viral titers. Taking advantage of an infectious SARS-CoV-2 strain expressing the chromophore mNeonGreen (14), we quantified infected (mNG+) and total (Hoechst+) cells by single-cell counting with an imaging multiplate reader. Of note, even short UV-C treatment of the dried virus in the context of the moving “fast” regimen completely inactivated SARS-CoV-2, as no infected cells were detected based on fluorescence protein expression (Fig. 1b). Titration of two-fold series dilutions of the UV-treated and non-treated control samples, as well as the freshly thawed strain as reference, revealed that (i) drying for two hours does not have a major impact on the infectivity of SARS-CoV-2 and (ii) all five UV-C treatment regimens effectively inactivate SARS-CoV-2 (Fig. 1c). Calculation of viral titers based on the titration of the reconstituted virus stocks revealed a loss of titer due to drying from ~4*10^6 to ~3*10^6 IU/ml in this set of experiments and effective 6-log titer reduction of SARS-CoV-2 by all employed UV-C treatment regimens down to 16 mJ/cm^2^ (Fig. 1d).

### Dose-dependent UV-C mediated inactivation of SARS-CoV-2

We next aimed to determine the UV-C doses at 254 nm sufficient to achieve complete disinfection respectively an at least 6-log reduction in viral titers. For this, we employed a battery-driven UV-C handheld device (Zone H) emitting 254 nm UV-C light at 0.7 mJ/cm^2^ at a distance of 50 cm. This allowed us to treat surface-dried SARS-CoV-2 with different UV-C doses by variation of the exposure time and additional use of a 97 % UV-C filter. In agreement with our previous measurement, drying for 2 hours did not significantly affect SARS-CoV-2 infectivity and relatively high doses of 254 nm UV-C treatment (14 mJ/cm^2^) inactivated SARS-CoV-2 (Fig. 2a exemplary images at 1:200 dilution and Fig. 2b quantitative analyses). Furthermore, there was a dose-dependent reduction in SARS-CoV-2 infectivity with total inactivation down to 3.5 mJ/cm^2^ while partial inactivation was still observed at 1.75 mJ/cm^2^ (Fig. 2a and b). Careful evaluation of viral titers post UV-C exposure revealed that > 6-log titer reduction was achieved by 3.5 mJ/cm^2^ 254 nm UV-C treatment (Fig. 2c). Of note, mean titers were only reduced by slightly more than one order of magnitude from 5.04*10^6^ IU/ml of the dried and reconstituted SARS-CoV-2 to 3.5*10^5^ IU/ml when the virus was exposed to 1.75 mJ/cm^2^, corresponding to 93 % inactivation. Therefore, the relationship between inactivation of surface dried SARS-CoV-2 and UV-C treatment is non-linear, at least in our system and 3.5 mJ/cm^2^ are necessary to achieve a 6-log titer reduction.

**Figure 2.**
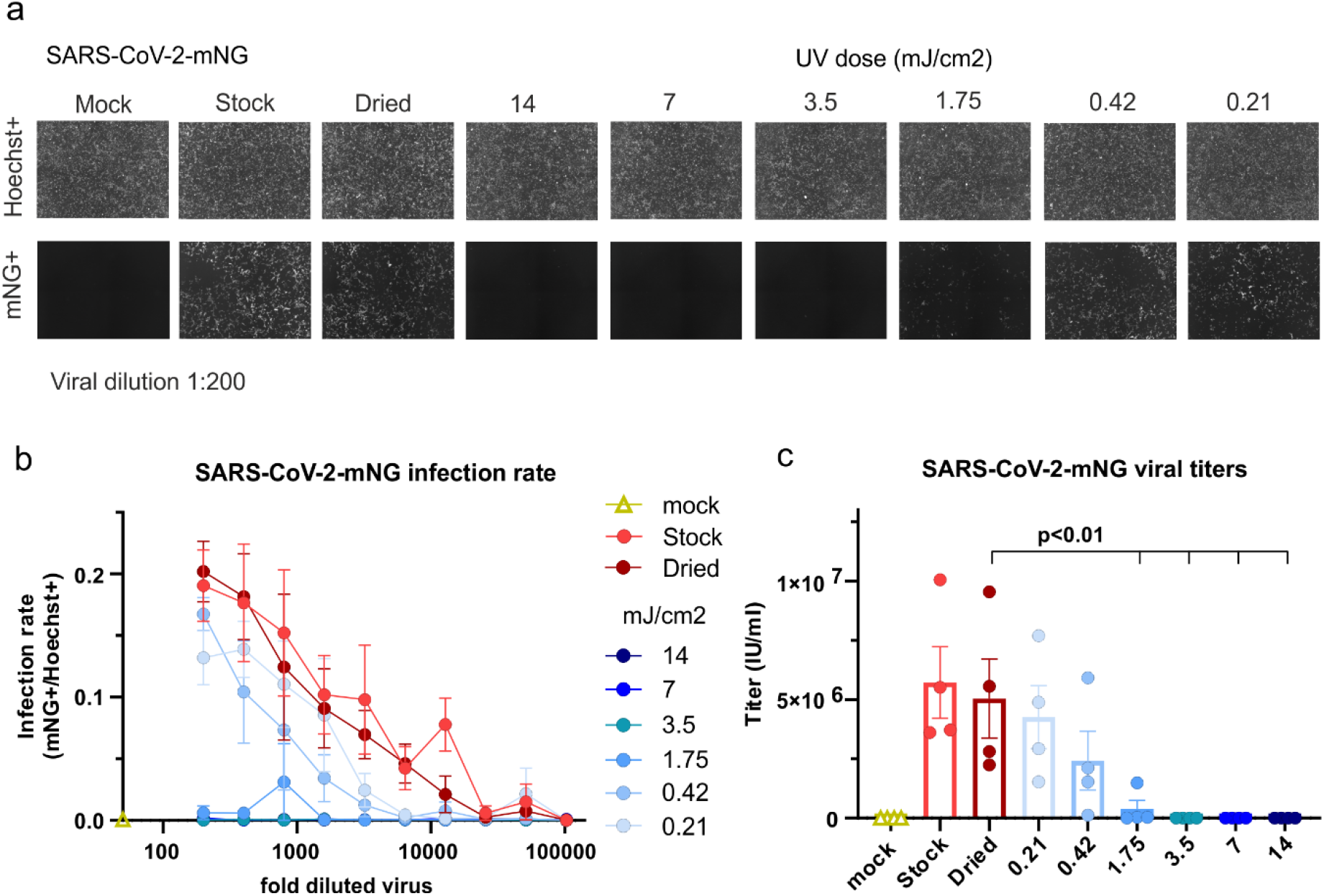
UV-C dose required for SARS-CoV-2 inactivation. (a) Primary data showing the results of the infection assay using mock-infected cells, non-treated stock virus as a positive control, and virus treated with the 6 UV-C doses as indicated. In the upper row, the total amount of cells is shown as Hoechst+. In the lower, infected cells at a viral dilution of 1:200 are visualized indicated as mNG+ cells. (b) Infection rate curves for UV-irradiated SARS-CoV-2-mNG using different UV-doses. The graph shows the infection rate at each two-fold serial dilution, calculated as the number of infected cells (mNG+) over the total number of cells (Hoechst+) for the non-treated viral stock, dried viral stock, and dried and UV-irradiated virus using different UV-C-doses (n=4). Data are presented as mean +/- SEM of the number of biological replicates indicated above. (c) SARS-CoV-2-mNG viral titers after UV-treatment. The graph shows the viral titers calculated in IU/mL for the mock-infected, non-treated, and dried stock as well as the dried and UV-irradiated virus under the different UV-C-doses. The number of biological replicates is n=4. Data are presented as mean +/- SEM. For analysis of statistical significance, we used a one-way ANOVA with multiple comparison and Fishers LSD-test.

## Discussion

Disinfection of surfaces and aerosols by UV-C irradiation is an established, safe and non-chemical procedure used for the environmental control of pathogens (1–3, 15). UV-C treatment has proven effective against several viruses including SARS-CoV-1 (4–6) and other coronaviruses i.e. Canine coronaviruses (16). Hence, as recently demonstrated by others (8–13) and now confirmed by our study it was expected that SARS-CoV-2 is permissive for inactivation by UV-C treatment. One critical question is the suitability of this technology in a setting in which the exposure time of surfaces or aerosols should be kept as short as possible to allow for a realistic application, for example in rooms that need to be used frequently as operating rooms or lecture halls. In such a setting, we assume that the virus is exhaled from an infected person by droplets and aerosols, dries on surfaces and hence represents a threat to non-infected individuals. We mimicked such a situation and first evaluated if surface dried SARS-CoV-2 is infectious. Drying for two hours, in agreement with previous work (13, 17), did not result in a significant reduction of viral infectivity indicating smear-infections could indeed play a role in the transmission of SARS-CoV-2 (Fig. 1). On the other hand, our virus-preparations are dried in cell culture pH-buffered medium containing FCS, which might stabilize viral particles. Hence, even though this is not the scope of the current study, it will be interesting to evaluate if longer drying or virus-preparations in PBS affect the environmental stability of SARS-CoV-2. Irrespective of the latter, UV-C-exposure of dried high-titer SARS-CoV-2 preparations containing ~3-5*10^6 IU/ml resulted in a complete reduction of viral infectivity (Fig. 1). In this context, it is noteworthy that we achieved a 6-log virus-titer reduction in a setting simulating surface disinfection with a moving handheld device. With the “fast”-moving protocol (see supplemental video 1) we were exposing surfaces at a distance of 20 cm with a speed of 12.5 cm/s resulting in a calculated integrated UV-C dose of 20 mJ/cm^2^ at 254 nm. This is substantially less than the previously reported 1048 mJ/cm^2^ necessary to achieve a 6-log reduction in virus titers when exposing aqueous SARS-CoV-2 to UV-C (10). In another study, using a 222 nm UV-LED source, 3 mJ/cm^2^ lead to a 2.51-log (99.7 %) reduction of infectious SARS-CoV-2 when irradiating for 30 s, however inactivation did not increase with extended irradiation regimens up to 300 s (9). In addition, 20 s deep-ultraviolet treatment at 280 nm corresponding to a dose of 75 mJ/cm^2^ reduced SARS-CoV-2 titer up to 3-logs (8). Finally, Storm and colleagues reported a 2-log reduction of dried SARS-CoV-2 at 4 s with 0.85 mW/cm^2^ corresponding to 3.4 mJ/cm^2^ (13). Of note, this value is highly similar to the dose of 3.5 mJ/cm^2^ calculated by us to be sufficient to achieve a > 6-log SARS-CoV-2 titer reduction, when the virus is in a dried surface residing state (Fig. 2). Comparing these values to other pathogens, SARS-CoV-2 seems particularly sensitive towards UV-C light. To achieve a 3-log titer reduction, 75-130 mJ/cm^2^ are necessary for adenovirus, 11-28 mJ/cm^2^ for poliovirus, and bacteria as for instance Bacillus subtilis require 18-61 mJ/cm^2^ (7).

Important limitations of UV-C-based disinfection procedures also exist. First and most importantly, UV-C irradiation is harmful to humans due to the high energy of the germicidal lamps and exposure of skin or eyes must be avoided. This excludes decontamination of populated public spaces by UV-C. Furthermore, UV-C does not penetrate surfaces, hence for efficient disinfection, equal direct irradiation of all surfaces with a sufficient dose has to be assured. Our work highlights this aspect, as due to the non-linear decay kinetic of the dose-response relationship 3.5 mJ/cm^2^ will totally inactivate high viral titers, whereas a slightly reduced dose of 1.75 mJ/cm^2^ only achieves roughly one-log reduction (Fig. 2c).

Apart from that, our study as well as the research done by others (13), emphasizes UV-C-based disinfection technologies as highly efficient to rapidly sterilize surfaces in different settings as for instance operating rooms, less-frequently populated areas in healthcare facilities and public transportation, but also in research facilities. Ideal applications are done in closed containers, precluding exposure of persons to UV-C radiation, when sterilizing small to medium-size objects. Another highly relevant aspect is the use of UV-C lamps in air sterilizers which would have a strong impact on public health and prevention of the public to infectious aerosols. However, the transferability of our results to viral aerosols, even though they give a first indicator, might be limited. Virus in aerosols exerts other dynamics and inactivation kinetics might differ. Hence, it is highly relevant and warranted to conduct studies to carefully determine UV-C doses necessary and sufficient for inactivation of SARS-CoV-2 in aerosols.

Altogether, we establish the effectiveness of UV-C treatment against SARS-CoV-2 in a setting designed to simulate close-to-reality conditions of decontamination. The easy, rapid, chemical-free, and high efficacy of UV-C treatment to inactivate SARS-CoV-2 demonstrates the applicability of this technology in a broad range of possible settings.

## Supporting information

SMovie_1

SMovie_2

Supplemental Images 1 and 2

## Author contributions

NR and MS designed the experiments; NR performed the experiments with support from RB; NR, RB and MS analyzed the data; NR and MS drafted the figures and wrote the manuscript; MS developed the manuscript to its final form; MS planned and supervised the study; all authors read, edited, and approved the final manuscript.

## Acknowledgements

We are thankful to Jan Winderlich, Jasmin Zahn, Christoph Söller, und Anika Hofmann (Heraeus) for fruitful discussions and for providing UV-C lamp spectra and UV-C dose emission measurements on the Soluva^®^ Zone H.

## Ethical statement

This study does not include any data obtained with primary patient cells or data. Hence, there was no necessity to obtain ethical approval by the internal review board.

**Supplemental Image 1. Spectrum of the UV-C lamps used.**

**Supplemental Image 2. UV-C emission at 254 nm of the Soluva^®^ Zone H at different distances and time points.**

**Supplemental Movie 1. UV-irradiation using the Handheld device, slow-moving regime.** SARS-CoV-2-mNG was spotted in a 6-well plate, dried for two hs and UV-irradiated as shown in the video. Speed is calculated at approx. 3.75 cm/s.

**Supplemental Movie 2. UV-irradiation using the Handheld device, fast-moving regime.** SARS-CoV-2-mNG was spotted in a 6-well plate, dried for two hs and UV-irradiated as shown in the video. Speed is calculated at approx. 12.5 cm/s.

