## Supplemental Images 1 and 2 for "Rapid, dose-dependent and efficient inactivation of surface dried SARS-CoV-2 by 254 nm UV-C irradiation"

This supplementary material is hosted by *Eurosurveillance* as supporting information alongside the article **“Rapid, dose-dependent and efficient inactivation of surface dried SARS-CoV-2 by 254 nm UV-C irradiation”**, on behalf of the authors, who remain responsible for the accuracy and appropriateness of the content. The same standards for ethics, copyright, attributions and permissions as for the article apply. Supplements are not edited by *Eurosurveillance* and the journal is not responsible for the maintenance of any links or email addresses provided therein.

**Supplemental Movie 1. UV-irradiation using the Handheld device, slow-moving regime.** SARS-CoV-2-mNG was spotted in a 6-well plate, dried for two hs and UV-irradiated as shown in the video. Speed is calculated at approx. 3.75 cm/s.

**Supplemental Movie 2. UV-irradiation using the Handheld device, fast-moving regime.** SARS-CoV-2-mNG was spotted in a 6-well plate, dried for two hs and UV-irradiated as shown in the video. Speed is calculated at approx. 12.5 cm/s.

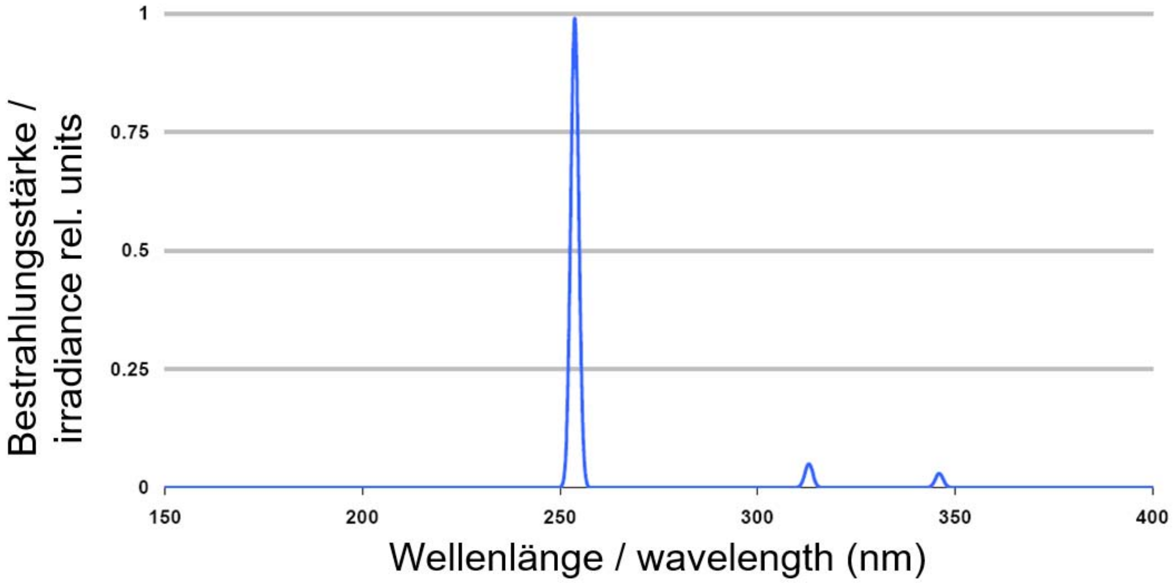


**Supplemental Image 1. Spectrum of the UV-C lamps used.**

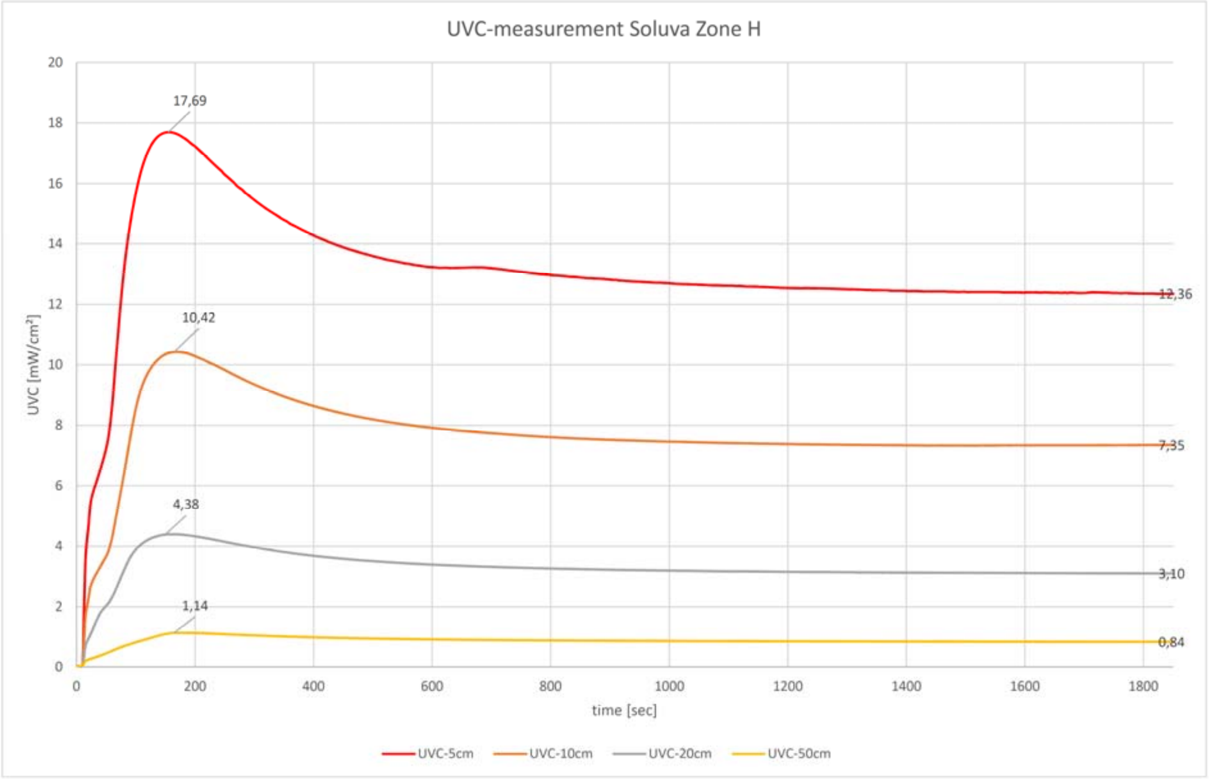


**Supplemental Image 2. UV-C emission at 254 nm of the Soluva® Zone H at different distances and time points.**
